# TAMC: A deep-learning approach to predict motif-centric transcriptional factor binding activity based on ATAC-seq profile

**DOI:** 10.1101/2022.02.15.480482

**Authors:** Tianqi Yang, Ricardo Henao

## Abstract

Determining transcriptional factor binding sites (TFBSs) is critical for understanding the molecular mechanisms regulating gene expression in different biological conditions. Biological assays designed to directly mapping TFBSs require large sample size and intensive resources. As an alternative, ATAC-seq assay is simple to conduct and provides genomic cleavage profiles that contain rich information for imputing TFBSs indirectly. Previous footprint-based tools are inheritably limited by the accuracy of their bias correction algorithms and the efficiency of their feature extraction models. Here we introduce TAMC (**<u>T</u>**ranscriptional factor binding prediction from **<u>A</u>**TAC-seq profile at **<u>M</u>**otif-predicted binding sites using **<u>C</u>**onvolutional neural networks), a deep-learning approach for predicting motif-centric TF binding activity from paired-end ATAC-seq data. TAMC does not require bias correction during signal processing. By leveraging a onedimensional convolutional neural network (1D-CNN) model, TAMC captures both footprint and non-footprint features at binding sites for each TF and outperforms existing footprinting tools in TFBS prediction particularly for ATAC-seq data with limited sequencing depth.

**AUTHOR SUMMARY:** Applications of deep-learning models are rapidly gaining popularity in recent biological studies because of their efficiency in analyzing non-linear patterns from feature-rich data. In this study, we developed a 1D-CNN model to predict TFBSs from ATAC-seq data. Compared to previous models using scoring functions and classical machine learning algorithms, our 1D-CNN model forgoes the need for bias correction during signal processing and significantly increases the efficiency in extracting features for TFBS prediction. In addition, the performance of our 1D-CNN model improves when the sequencing depth of training ATAC-seq data increases. Importantly, we showed that our method outperforms existing tools in TFBS prediction particularly when the sequencing depth of training ATAC-seq data is higher than the ATAC-seq data for prediction. This widened the applicability of our model to ATAC-seq data with both deep and shallow sequencing depth. Based on these results, we discussed about the potential application of our method to TFBS predication using bulk and single-cell ATAC-seq data.

## INTRODUCTION

Transcription factors (TFs) are proteins that bind to conserved genomic sequence motifs and have functions in regulating gene expression (1). Determining TF binding sites (TFBSs) is essential for deciphering the molecular mechanisms regulating gene expression across different biological conditions. Biological assays, such as ChIP-seq (2) and CUT&RUN (3), have been used as the standard experimental methods for mapping genome-wide interactions between TFs and chromatin. However, these experiments are resource-intensive and can measure only one TF at one time, which largely limits their applications in many situations. To address these limitations, computational methods, discussed below, have been proposed to impute TFBSs.

Traditional computational TFBS prediction methods have been using position weight matrices (PWMs) of TF binding motifs against DNA sequence to generate motif-predicted binding sites (MPBSs) (4, 5). However, all these MPBS-based methods suffer from high false-positive rates (FDR) (6). Recent studies have shown that more than 90% of TF binding events take place at open chromatin regions (7) that could be mapped by enzymatic cleavage assays such as DNase-I sequencing (DNase-seq) (8) and Assay for Transposase Accessible Chromatin sequencing (ATAC-seq) (7). Notably, bound TFs hinder the activity of cutting enzymes and leave footprint sites characterized with lower cleavage signal frequency comparing to surrounding regions (9). Therefore, the TF-bound and unbound MPBSs can be theoretically distinguished by their footprint pattern.

Several computational methods have been developed to investigate footprint patterns in chromatin cleavage profiles (10-19). As chromatin cleavage events generated by cutting enzymes (e.g., DNase-I used in DNase-seq and Tn5 transposase used in ATAC-seq) are biased towards different sequences, previous tools initially designed for DNase-seq data often give poor predictions using ATAC-seq data (16, 20, 21). By far, HINT-ATAC (16) and TOBIAS (17) are two representative footprinting tools specifically designed for ATAC-seq data, which has become the dominant data type for chromatin accessibility profile because of the simplicity of the assay itself.

TOBIAS uses a simple footprint score (FPS) metric to characterize the footprint pattern at single-base resolution while HINT-ATAC uses a semi-supervised hidden Markov model (HMM) to predict footprint sites directly. Both tools significantly increase the accuracy in classifying bound/unbound MPBSs from ATAC-seq data. On the other hand, the limitation is that they both require complex bias correction during signal processing because their models/algorithms are highly dependent on measurable footprint pattern to make predictions.

More recently, deep-learning models are rapidly gaining popularity in biological studies because of their efficiency in analyzing complex (non-linear) patterns from feature-rich data. Here, we introduced a new TFBS prediction tool named TAMC (**<u>T</u>**ranscriptional factor binding prediction from **<u>A</u>**TAC-seq profile at **<u>M</u>**otif-predicted binding sites using **<u>C</u>**onvolutional neural networks). TAMC takes advantage of signal processing strategies in HINT-ATAC and TOBIAS except for the bias correction step to produce input signals that are then used to extract features of TFBSs using a 1D-convolutional neural network (1D-CNN) model (Figure 1A). By evaluating TAMC with different input signal configurations, we showed that TAMC does not require bias correction during signal processing and captures both footprint and non-footprint features of TFBSs efficiently. Importantly, TAMC pretrained with multiple deeply sequenced ATAC-seq datasets significantly outperforms HINT-ATAC and TOBIAS under both intra-data and cross-data settings. This makes TAMC a competitive alternative method for TFBS prediction especially for ATAC-seq data with limited sequencing depth.

**Figure 1.**
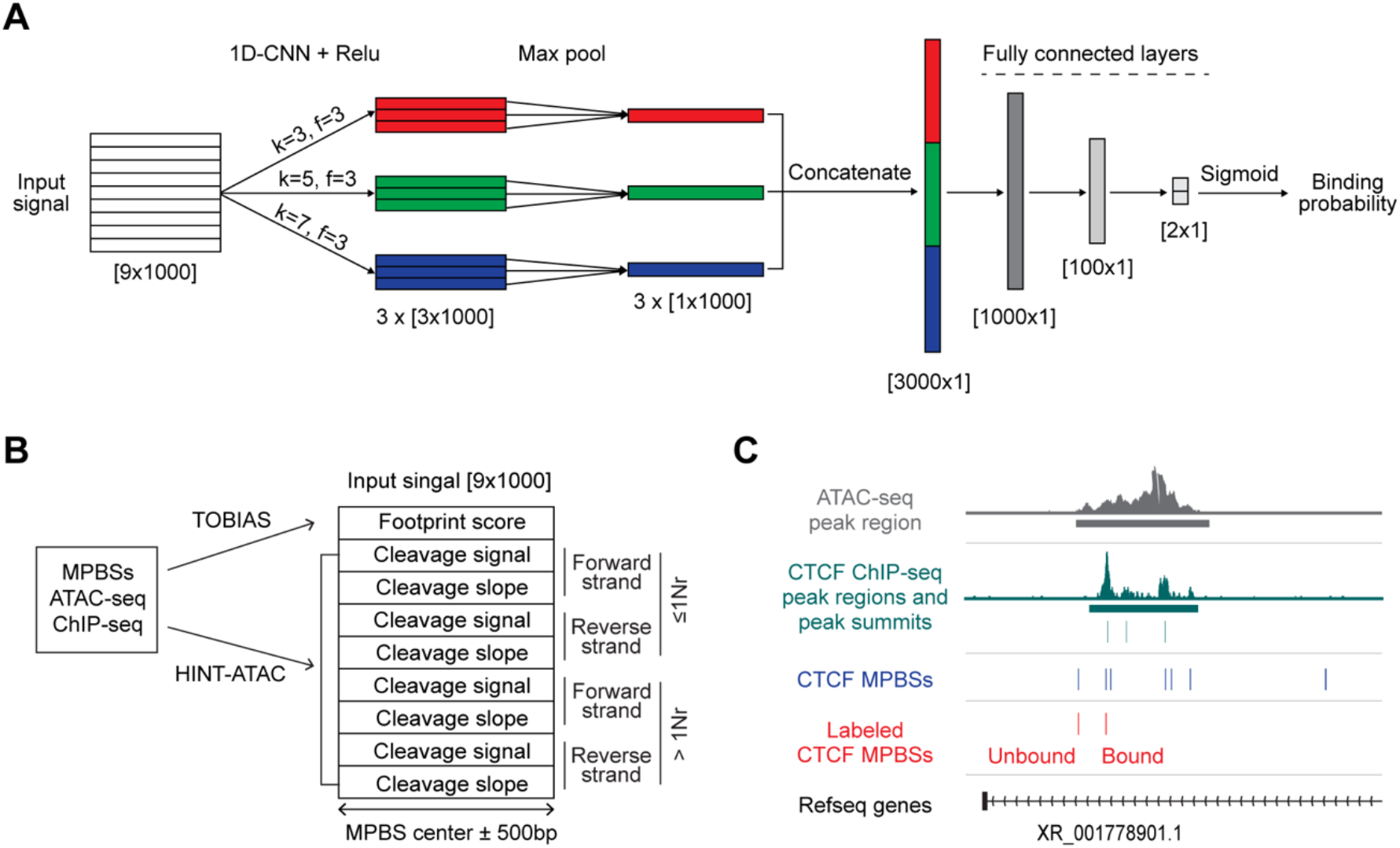
TAMC model and input signal. (A) Architecture of TAMC framework: three convolutional layers with 3 learnable filters (f=3) of kernel sizes k=3, 5, and 7, and Relu activations, followed by max pooling, concatenation, and prediction of binding probability via a two fully connected layers with sigmoid activation. (B) Default TAMC input signal structure. >1Nr, ATAC-seq read fragments larger than one nucleosome size; ≤1Nr, ATAC-seq read fragments equal or smaller than one nucleosome size. (C) Representative tracks show strategy of labeling bound and unbound CTCF MPBSs in GM12878 cell type.

## RESULTS

### TAMC overview

TAMC takes a combination of footprinting scores and genomic cleavage profile (signals and slopes) around 500bp from the center of each MPBS as input signal (Figure 1B). The footprint scores are calculated using the TOBIAS footprint scores metric, and the genomic cleavage signals and slopes are generated following HINT-ATAC input signal processing strategy except for the bias correction step (Figure 1B). The genomic cleavage signals and slopes are further separated into 8 channels by strand and size of ATAC-seq reads and fragments (Figure 1B). Finally, the resulting 9-channel input signals are fed into a 1D-CNN module using three kernel sizes (k=3, 5, and 7), each of which has 3 learnable kernels to extract high-level features at different resolutions (Figure 1A). The extracted feature maps are then aggregated using max pooling (i.e., extracting the most salient feature from the 3 filters at each resolution, k) and concatenated before being fed to the three fully connected layers and a final sigmoid activation function to make predictions of TF binding probability (Figure 1A).

The TAMC framework was trained and tested using published paired-end ATAC-seq data of three human cell types (GM12878, HepG2 and K562) (Table S1). ChIP-seq data obtained from the same cell type as ATAC-seq data were used for labeling each MPBS as bound or unbound (Figure 1C). Because of the high FDR of MPBSs (6), there are far more unbound MPBSs than bound MPBSs (Figure S1A). To avoid biases due to training and evaluation with a disproportionally large unbound category, we randomly subsampled the bound and unbound MPBSs to have equal proportions for each TF before conforming training, validation and testing datasets (Figure S1B). The resulting balanced datasets are large enough (more than 5000 labeled MPBSs for most TFs) to enable reliable model training and performance metrics robust to the random subsampling. The trained TAMC models were tested under two types of prediction settings: intra-data prediction (same ATAC-seq data for training and testing) and cross-data prediction (different ATAC-seq data sets for training and testing). In total, we trained and tested TAMC for 47 TFs with published ChIP-seq data for all three cell types in the ENCODE database (Table S2). Area under the receiver operating curve (AUC) was used to measure the trained models’ performance in classifying bound/unbound MPBSs.

### TAMC outperforms existing methods

To evaluate TAMC performance, we compared TAMC predictions with predictions generated using previous footprinting tools including TOBIAS and HINT-ATAC. We showed that TAMC gave the best intra-data predictions for most of the 47 TFs in both GM12878 and HepG2 cells (Figure 2A). To get cross-data prediction performance of TAMC, we trained TAMC using GM12878 and/or HepG2 ATAC-seq data and then tested the trained models using ATAC-seq data of another cell type, K562. Such cross-data training and testing were conducted using ATAC-seq datasets downsized to different total aligned reads number in order to check the influence of sequencing depth on TAMC performance. Our results showed that TAMC models trained using multiple ATAC-seq datasets gave better cross-data predictions than models trained using a single ATAC-seq data (Figure 2B). In addition, the higher the sequencing depth of the training datasets is, the better cross-cell prediction performance could be provided by TAMC (Figure 2C). Compared to TOBIAS and HINT-ATAC, TAMC gave best cross-cell prediction when the sequencing depth of testing datasets is low; while for testing data with high sequencing depth, TOBIAS gave better prediction than TMAC (Figure 2C). Interestingly, the performance of TOBIAS and TAMC were always crossed near the point where the training and testing ATAC-seq datasets have similar sequencing depth (Figure 2C). These results together suggested that TAMC outperforms TOBIAS and HINT-ATAC in cross-data TFBS prediction as long as the training ATAC-seq datasets were sequenced at higher depth than the testing ATAC-seq datasets.

**Figure 2.**
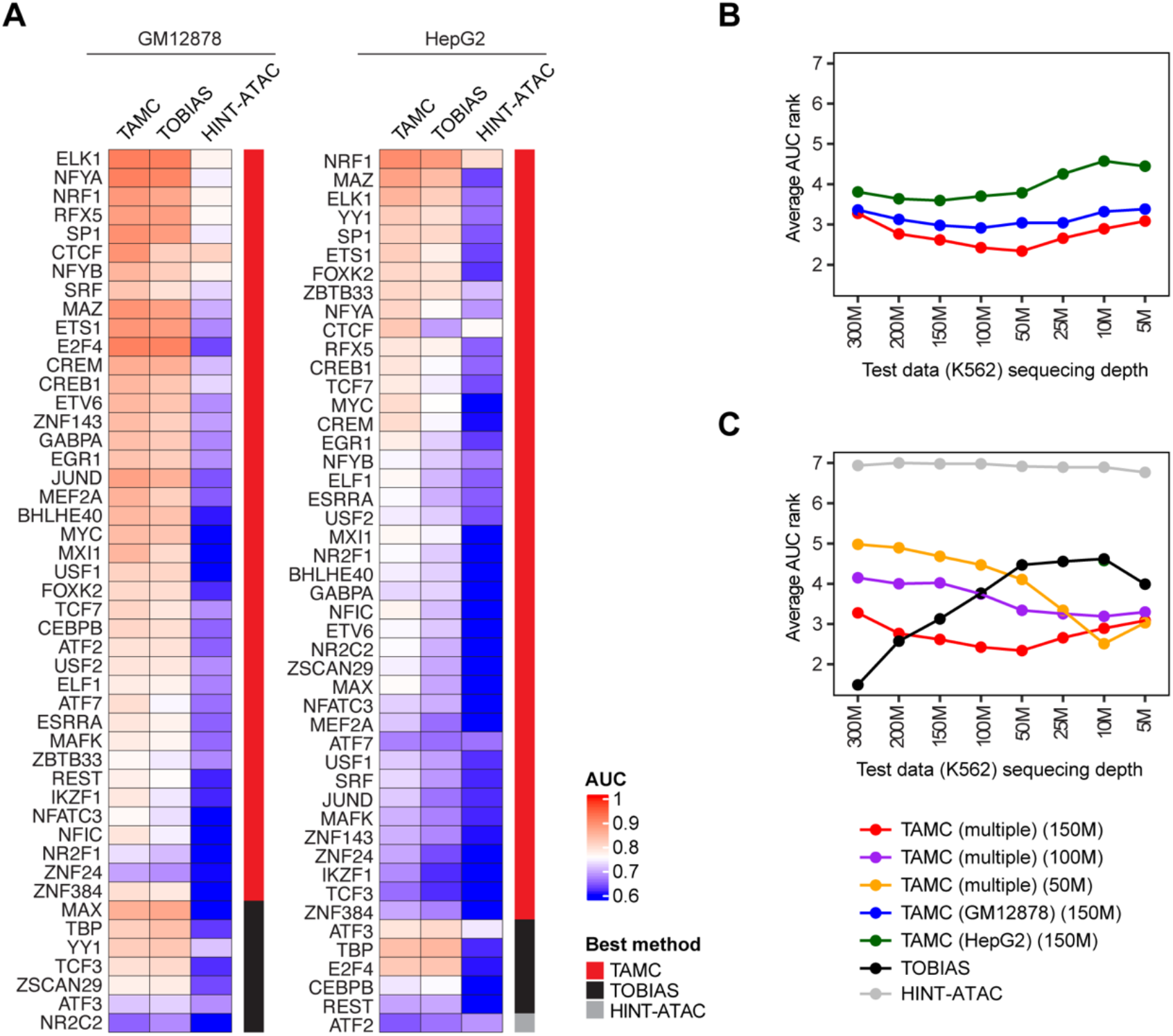
TAMC outperforms existed methods. (A) Heat map compares intra-data prediction performance of TAMC, TOBIAS and HINT-ATAC cells for each TF in GM12878 and HepG2. The method giving best prediction for specific TF is indicated in the side bar. (B-C) Line graphs compare average cross-data prediction performance of TAMC, TOBIAS and HINT-ATAC for all 47 TFs. TAMC models were trained using GM12878, HepG2 or multiple (GM12878 + HepG2) ATAC-seq datasets. The cell type and sequencing depth of ATAC-seq datasets used for training TAMC models were marked within parentheses. All models were tested using K562 ATAC-seq data downsized to 8 different sequencing depths. The cross-data performance of all 7 models were ranked from 1 to 7 for each TF. The higher AUC is, the lower rank number is given. M denotes million reads.

### TAMC does not require bias correction during input signal processing

Most existing footprinting tools, including TOBIAS and HINT-ATAC, require conducting bias correction during cleavage signal processing to uncover measurable footprint patterns for bound/unbound MPBS classification. However, none of the reported bias correction algorithms can uncover footprint for all TFs faithfully (17). This makes bias correction a critical step that limits the performance of existing footprint-based methods.

By default, TAMC does not conduct bias correction during signal processing. To check whether TAMC performance is affected by bias correction as most footprinting methods do, we compared the performance of TAMC models using four different combinations of bias-corrected and uncorrected signals as inputs together with TOBIAS and HINT-ATAC (Figure 3). Our results showed that all four TAMC models gave better intra-data predication results than TOBIAS and HINT-ATAC in both GM12878 and HepG2 cells (Figure 3). However, we did not observe consistent difference in the performance between the four TAMC models in the two cell types (Figure 3). Therefore, TAMC can distinguish between bound and unbound MPBS without the requirement of bias correction during input signal processing.

**Figure 3.**
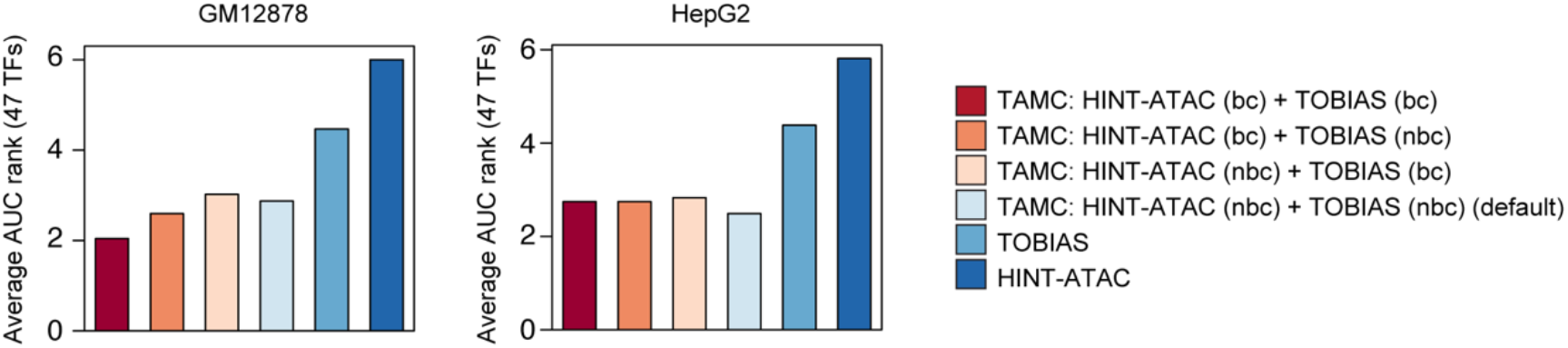
TAMC performance is independent of bias correction. Bar graphs compare average intra-data prediction performance for 47 TFs using TOBIAS, HINT-ATAC and TAMC models with four different combinations of bias-corrected and uncorrected signals as inputs in GM12878 and HepG2 cells. The higher AUC is, the lower the rank number. bc, bias corrected; nbc, non-bias corrected.

### TAMC generates TF-specific models

Both TOBIAS and HINT-ATAC assume that the footprints left by all TFs have the same pattern across the genome – TOBIAS applies the same matric to calculate the footprint scores at all genomic sites and HINT-ATAC uses the model trained for EGR1 to make predictions for all TFs. However, it has been recently reported that the footprint pattern for different TFs are highly heterogeneous from each other (10). To check whether TAMC can tell the heterogeneity in binding features of different TFs, we compared intra-data prediction performance of TAMC using models trained for the same or different TFs as the testing TF. We determined 6 TFs (ZNF143, NR2F1, CTCF, MXI1, NFIC and MEF2A) always require TAMC models trained using the same TF for best binding site prediction (Figure 4). In particular, the model trained for CTCF showed extremely high specificity for CTCF binding site prediction (Figure 4). This result suggested that TAMC can tell TF-to-TF differences at their binding sites and thus generate TF-specific models to improve prediction accuracy.

**Figure 4.**
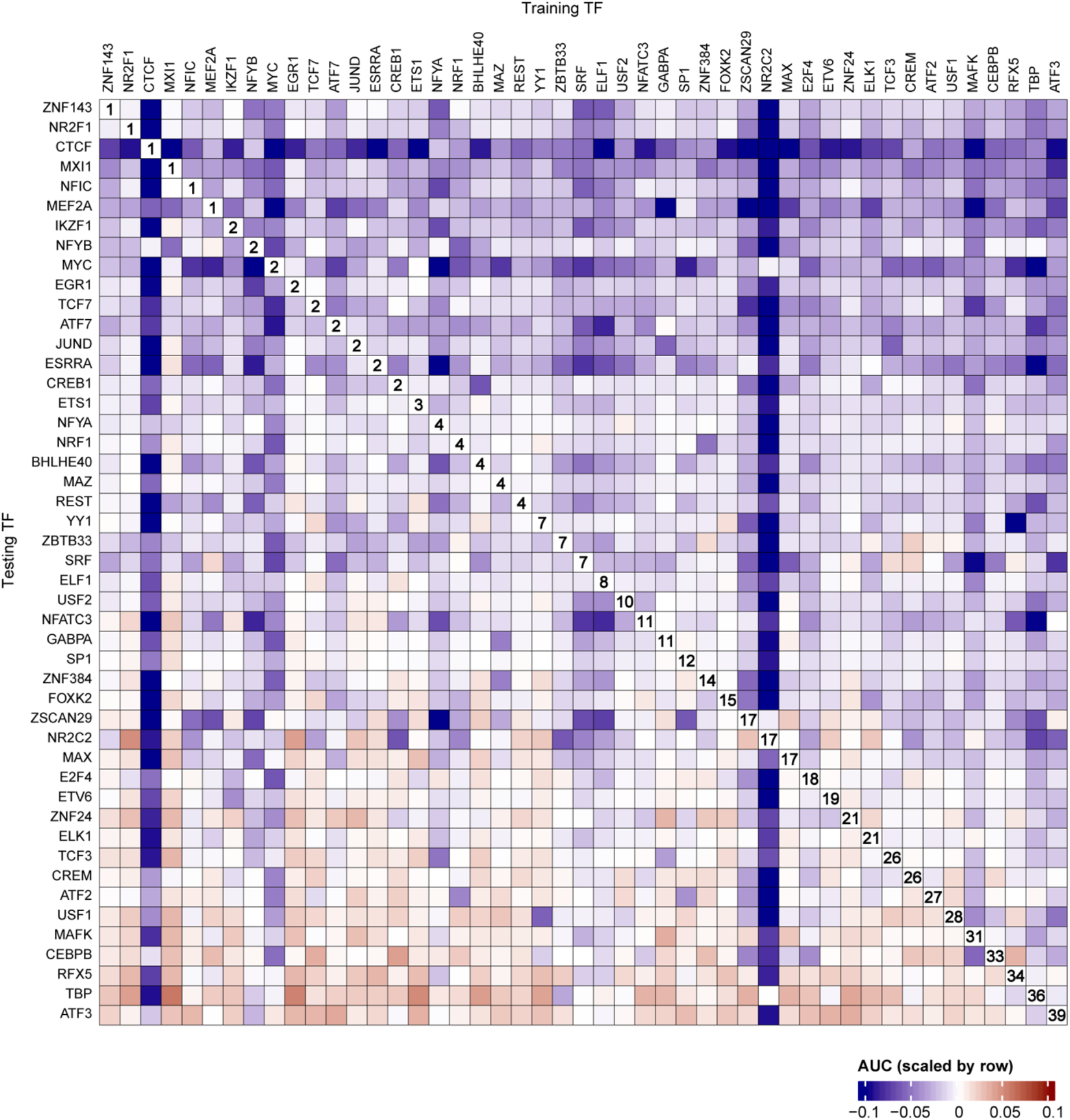
TAMC generates TF-specific models. Heat map compares intra-data prediction performance of TAMC using models trained for the same or different TFs as the testing TF in GM12878 cell. For each testing TF, AUC values of TAMC models trained for 47 TFs were scaled between the range [-0.1, 0.1]. The ranks for models trained with the same TF as the testing TF is labeled in the plot.

### TAMC captures both footprint and non-footprint features of TFBSs by deep learning

To further explore how TAMC exceeds TOBIAS and HINT-ATAC in predicting TF binding dynamics at MPBSs, we tested TAMC with four variant input structures that lack footprint scores (variant 1), cleavage profile (variant 2), read strand (variant 3) and fragment size (variant 4) information respectively (Figure 5A).

**Figure 5.**
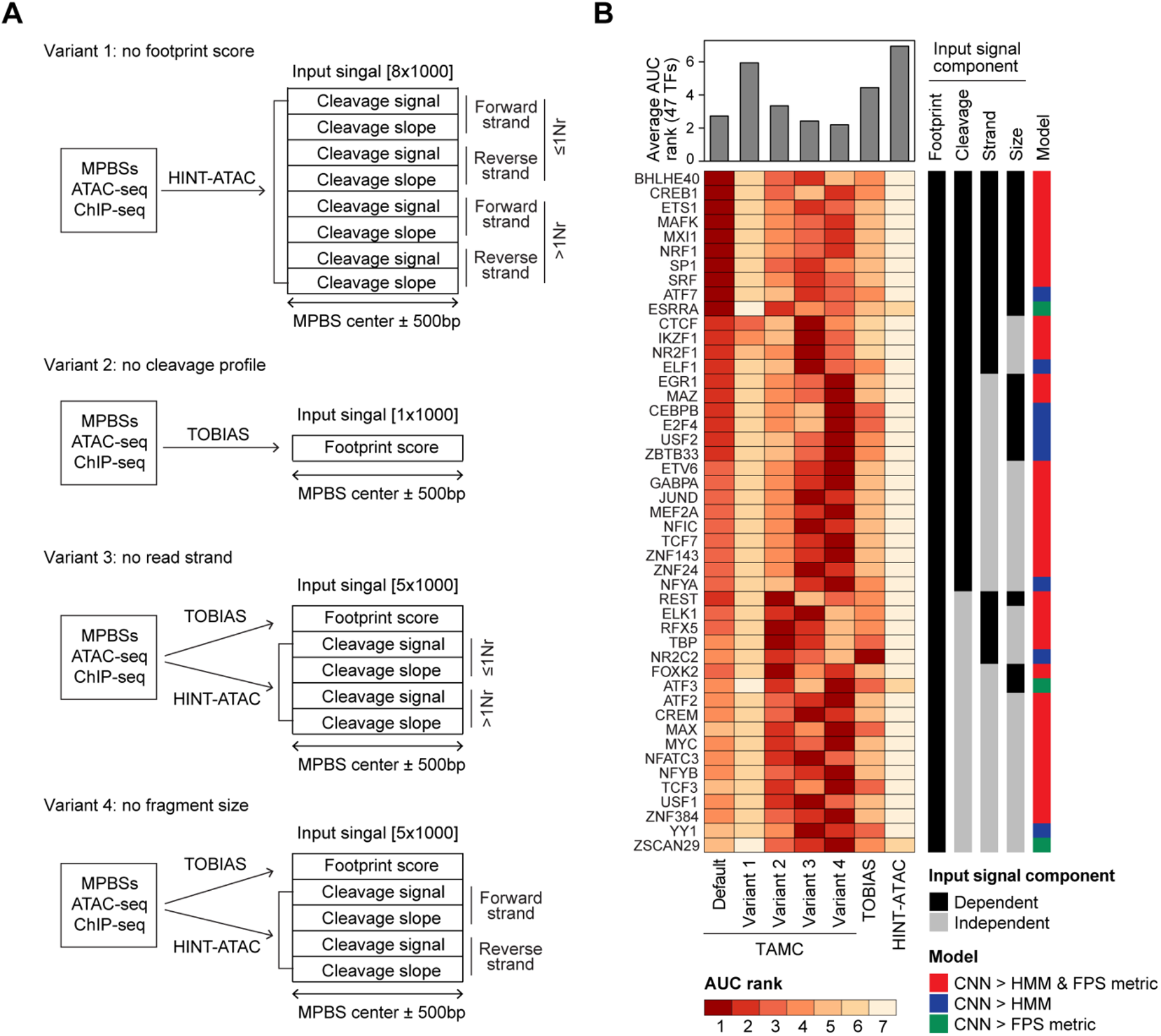
TAMC captures both footprint and non-footprint features of TFBSs by deep learning. (A) Variant TAMC input structures that lack footprint score, cleavage profile, strand and size information respectively. (B) Heat map compares intra-data prediction performance of TAMC using default and variant input structures together with TOBIAS and HINT-ATAC in GM12878 cell. The performance the 7 models were ranked from 1 to 7 for each TF. TFs that require footprint score, cleavage profile, strand or size information within the input signal or the 1D-CNN model for better prediction are indicated in the side bar.

By comparing intra-data prediction performance of default and variant TAMC models, we can define what kind of information in the input signals are important for TAMC to make predictions. Our results showed that TAMC performance was drastically compromised for all 47 TFs when footprint score is removed from input (Figure 5B). In addition, loss of cleavage profiles completely impaired TAMC prediction accuracy for more than half of the TFs (Figure 5B). Specifically, we determined several TFs requiring some non-footprint information that are only provided by the cleavage profile part, including read stand (e.g., CTCF) and/or fragment size (e.g., EGR) information, for better binding site prediction (Figure 5B). Therefore, both the footprint score part and cleavage profile part in TAMC input signal can provide important information for characterizing TFBSs.

At the same time, by comparing TAMC models using variant 1 and 2 input structures with HINT-ATAC and TOBIAS respectively, we showed that the 1D-CNN model used in TAMC makes better predictions than the classical model (e.g., HMM in HINT-ATAC) or non-model metrics (e.g., FPS metric in TOBIAS) even using the same input signals (Figure 5B). Therefore, the high efficiency in capturing complex and subtle features by 1D-CNN model also played an important role in improving TFBS prediction accuracy by TAMC.

## DISCUSSION

Predicting TF binding sites is important for understanding gene expression mechanisms. In this study, we introduced a new tool named TAMC to predict TF binding dynamics at MPBSs using paired-end ATAC-seq data. As summarized in Table 1, TAMC has several advantages comparing to previous tools. First, TAMC does not require bias correction during signal processing, which makes signaling processing easier and avoids further artificial bias caused by the bias correction algorithm. Second, TAMC combines different configurations of processed ATAC-seq signals within its input and therefore retains not only footprint but also non-footprint information, such as stand and size information, for later prediction use. Finally, TAMC uses 1D-CNN to analyze input signals, which is based on our results, more efficient than classical models (e.g., HMM in HINT-ATAC) or non-model-based matrices (e.g., FPS matric in TOBIAS) in feature capturing. These advantages together allow TAMC to capture precise binding features of each TF and thus improves its performance in classifying bound/unbound MPBSs.

**Table 1.** Summary of features of TAMC and existing footprinting tools for ATAC-seq data.

| **Tools** | **TAMC** | **TOBIAS** | **HINT-ATAC** |
| --- | --- | --- | --- |
| **Year of publication** | This paper | 2020 | 2019 |
| **Bias correction** | Not required | Required | Required |
| **Input** | Footprint score +<br>Cleavage profile | Footprint score | Cleavage profile |
| **Model** | 1D-CNN | FPS metric | HMM |
| **Feature captured for**<br>**TFBS prediction** | Footprint +<br>Non-footprint | Footprint | Footprint +<br>Non-footprint |
| **Performance**<br>**evaluation setting** | Intra-data +<br>Cross-data | Intra-data | Intra-data |

Most previous studies use the training and testing datasets derived from the same ATAC-seq data for evaluating the tools’ performance in classifying bound/unbound MPBSs. However, in real applications, the cell type and sequencing depth of ATAC-seq data for prediction are usually different from available training data. To mimic the situations in real application, we evaluated TAMC performance under both intra-data and cross-data settings. We showed that TAMC outperforms existed methods as long as the training datasets has higher sequencing depth than testing datasets. In real studies, bulk ATAC-seq assays are usually conducted at sequencing depth between 100~200M reads and can provide 20~50M high-quality aligned reads after removing duplicated and mitochondrial DNA. In our study, the TAMC models are trained using ATAC-seq data with around 150 million high-quality aligned reads. Therefore, the trained TAMC models generated in this study are suitable for TFBS prediction using most bulk ATAC-seq data generated in real studies. Recently single-cell ATAC-seq is becoming more and more popular because it can analyze multiple cell types at one time. One major obstacle for TFBS prediction using scATAC-seq data is its low sequencing depth for individual cell or specific cell population. As our TAMC models significantly exceeded existing methods in TFBS prediction using ATAC-seq data with low sequencing depth, it could be further applied in analyzing TF binding dynamics using scATAC-seq data in the future.

Among the 47 TFs analyzed in our study, only a small number of TFs (e.g., ATF3 and TBP) did not show better binding site prediction result by TAMC comparing to existed methods (Figure 2A). We noted that these TFs often have very low number of bound MPBSs (Figure S1), which could result in insufficient data amount for model training. One possible way to improve TAMC’s performance for these TFs is to combine labeled MPBSs from different cell types for training. Importantly, our results revealed that in addition to increasing data amount, combining training datasets of multiple cell types can prevent cell type-specific overfitting effect as well, which is important for making cross-data predictions in real applications (Figure 2B). Therefore, while the trained TAMC models generated in our study already outperformed TOBIAS and HINT-ATAC for most TFs, more robust and generalized models could be obtained by combining deeply sequenced ATAC-seq data of more cell types for training.

## METHODS

### ATAC-seq and ChIP-seq data processing

ATAC-seq and ChIP-seq data (Tables S1 and S2) were obtained from the UCSC ENCODE portal (https://www.genome.ucsc.edu/ENCODE). Raw ATAC-seq and ChIP-seq fastq files were trimmed using TrimGalore (22) and aligned with Bowtie2 (v2.3.5.1, --qc-filter --very-sensitive) (23) to reference human genome (h38). Reads with alignment quality lower than 30 or reads aligned to mitochondrial DNA were removed using Samtools (v1.10) (24). Duplicated reads were removed using MarkDuplicates tool of Picard (v2.0.1; http://broadinstitute.github.io/picard/). The aligned bam files of GM12878 and HepG2 ATAC-seq datasets were further downsized to 150, 100 and 50 million total reads before used for training TAMC models. The aligned bam file for K562 ATAC-seq data was also downsized to 300, 200, 150, 100, 50, 25, 10 and 5 million aligned reads for testing trained TAMC models under different sequencing depth situations. Both ATAC-seq and ChIP-seq peaks regions and summits were called using MACS2 (v2.2.7.1, --nomodel --nolambda --keep-dup auto --call-summits) (25).

### MPBS detection and labeling

Binding motifs for 47 TFs were all obtained from JASPAR CORE 2020 database (https://jaspar.genereg.net/). For TFs with redundant motifs, only the latest version of motifs was used. MPBSs for each TF across hg38 genome were mapped using the MOODS with a p-value threshold of 0.0001 (4). Only MPBSs located within open chromatin regions (ATAC-seq peak regions) were used for further analyses. To label the TF-bound and unbound status of each MPBS, ChIP-seq data obtained from the same cell type as ATAC-seq data were used. MPBSs located outside ChIP-seq peak regions were labeled as unbound sites. For MPBSs located within each ChIP-seq peak region, only the MPBS located closest to and within 50bp from the highest ChIP-seq summit within that peak was kept and labeled as TF-bound in later analyses. If one TF has more unbound MPBSs than bound MPBSs, the obtained unbound MPBSs will be further randomly downsized to the same number as bound MPBSs for further analyses and vice versa.

### Input signal processing

To prepare input signals for TAMC, the ATAC-seq data was processed using ATACorrect and FootprintScores tools in TOBIAS package to generate footprint scores at single-based resolution (17). Footprint scores within 500bp form the center of each MPBS was made into a 1×1000 footprint feature vector. At the same time, the ATAC-seq data were separated into 4 files by strand and fragment sizes, and then each file was used for counting genomic cleavage signals and calculating slopes of cleavage signals within 500bp from each MPBS center following the signal processing steps in HINT-ATAC package (16). The obtained 4-channels of genomic cleavage signals and 4 channels of cleavage slopes were combined into 8×1000 cleavage feature vectors. The default TAMC input signal was generated by concatenating the 1×100 footprint feature vector and the 8×1000 cleavage feature vector to form the 9×1000 input feature vector. In addition, the 1×1000 footprint feature vector and the 8×1000 cleavage feature vector form two variant TAMC input structures lacking cleavage profiles and footprint information by themselves. To make the 5×1000 variant input feature vectors lacking size or strand information, the 1×1000 footprint feature vector was concatenated with a 4×1000 cleavage feature vector generated using ATAC-seq data only separated by strand or fragment sizes respectively.

### Training

The TAMC model was trained using input signals generated using ATAC-seq data of GM12878 and/or HepG2 cells. For each TF, 70% of labeled MPBSs were used for training and 20% for validation. The bound and unbound MPBSs for each TF should have equal number in training and validation datasets to avoid biased training. The python package PyTorch (26) was used for generating and training the model (Figure 1A). The model is trained using the Adam algorithm with minibatch size of 35, epoch size of 10 and maximum iteration number of 100. The datasets were randomized between each epoch of training. Validation loss is evaluated at the end of each training iteration and trained models with lowest 3 validation losses were saved as best models.

### Evaluation

The trained TAMC models were tested sing ATAC-seq data of GM12878, HepG2 or K562 cells. For GM12878 and HepG2 cells, the remaining 10% labeled MPBS were used to prepare testing input signals. For K562 ATAC-seq data, all labeled MPBS were used for testing. The bound and unbound MPBSs in testing datasets for each TF have equal proportions to avoid a biased evaluation. For each TF, the AUC value was calculated based performance in classifying bound/unbound MPBS using the average of binding probabilities predicted by the 3 best models.

The performance of TAMC was compared with TOBIAS and HINT-ATAC. To evaluate TOBIAS performance, the AUC was calculated based on the ability of the average footprint scores within each MPBS to classify bound/unbound sites. To evaluate HINT-ATAC performance, all MPBSs were first rank by the PWM scores generated by MOODS and the maximum PWM score was saved. Then footprint regions were identified using the footprinting tool of rgt-hint package (16). MPBSs overlapping footprint regions were assumed to be bound and their PWM scores were renewed by adding the maximum PWM score. The other MPBSs were regarded as unbound and their PWM scores were kept unchanged. The AUC is calculated based on the ability of the final PWM scores in classifying bound/unbound MPBSs.

## Code availability

TAMC scripts for input signal preparation, training and making predictions are publicly available at GitHub (https://github.com/tianqiyy/TAMC).

## Acknowledgements

We thank Dr. Lawrence Carin for providing suggestions for this project and Shelley Rusincovitch for organizing the Duke Data Science Plus (+DS) program. For project involving non-sensitive data, +DS is supported by Duke Research Computing for the use of the Duke Compute Cluster for high-throughput computation. The Duke Data Commons storage is supported by the National Institutes of Health (1S10OD018164-01).

## SUPPLMENTARY MATERIAL

**Figure S1.**
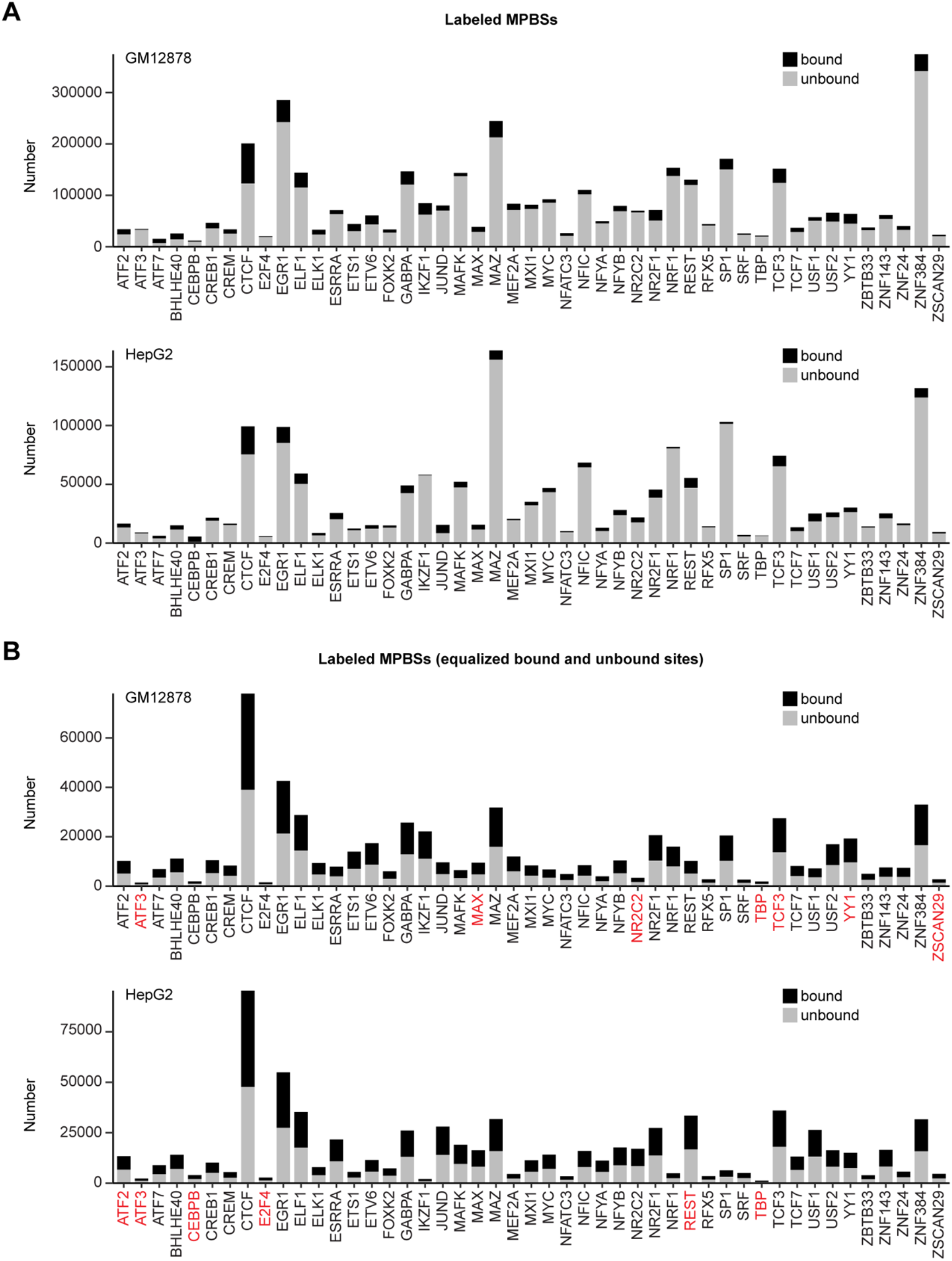
Number of labeled MPBSs for 47 TFs before (A) and after (B) equalization of bound and unbound sites in GM12878 and HepG2 cell lines. TFs did not show better binding sites prediction using TAMC than TOBIAS or HINT-ATAC were highlighted in red.

**Table S1.** List of ATAC-seq datasets.

| Cell line | ENCODE datasets | ENCODE data accession | Total aligned reads number |
| --- | --- | --- | --- |
| GM12878 | ENCSR637XSC | ENCFF063GWP<br>ENCFF095PJX<br>ENCFF948NNH<br>ENCFF181DBZ<br>ENCFF747WKN<br>ENCFF225QOQ | 155,940,597 |
| HepG2 | ENCSR291GJU | ENCFF875IIN<br>ENCFF461JXR<br>ENCFF134LRP<br>ENCFF734TQO<br>ENCFF483VIK<br>ENCFF039OZA | 152,034,393 |
| K562 | ENCSR868FGK | ENCFF261UXP<br>ENCFF123TMX<br>ENCFF098UCE<br>ENCFF703BGR<br>ENCFF902AAO<br>ENCFF178ILL<br>ENCFF431UDN<br>ENCFF147WEL<br>ENCFF774CPY<br>ENCFF160WRX<br>ENCFF354EXH<br>ENCFF575ZTZ<br>ENCFF260KVA<br>ENCFF761EDP<br>ENCFF819HSY<br>ENCFF168MQE<br>ENCFF457HSH<br>ENCFF441SZZ<br>ENCFF962SRP<br>ENCFF420XJE<br>ENCFF903FIK<br>ENCFF924YGE.<br>ENCFF827WXQ<br>ENCFF742FCG | 531,852,489 |

**Table S2.** List of TF motifs and ChIP-seq datasets.

| TF | Motif<br>(JASPAR 2020) | ENCODE datasets |  |  |
| --- | --- | --- | --- | --- |
|  |  | GM12878 | HepG2 | K562 |
| ATF2 | MA1632.1 | ENCSR000BQK | ENCSR047BUZ | ENCSR869IUD |
| ATF3 | MA0605.2 | ENCSR000BJY | ENCSR000BKE | ENCSR028UIU |
| ATF7 | MA0834.1 | ENCSR014YCR | ENCSR545FXC | ENCSR972ZBV |
| BHLHE40 | MA0464.2 | ENCSR000DZJ | ENCSR000EDT | ENCSR000EGV |
| CEBPB | MA0466.2 | ENCSR000BRX | ENCSR000EEE | ENCSR000EHE |
| CREB1 | MA0018.3 | ENCSR000BUF | ENCSR000BVL | ENCSR000BSO |
| CREM | MA0609.2 | ENCSR839XZU | ENCSR903ELW | ENCSR077DKV |
| CTCF | MA0139.1 | ENCSR000DZN | ENCSR000BIE | ENCSR000EGM |
| E2F4 | MA0470.2 | ENCSR000DYY | ENCSR924LSO | ENCSR000EWL |
| EGR1 | MA0162.4 | ENCSR000BMQ | ENCSR026GSW | ENCSR024CNP |
| ELF1 | MA0473.3 | ENCSR000BMB | ENCSR000BMZ | ENCSR502OEK |
| ELK1 | MA0028.2 | ENCSR000DZB | ENCSR717QSS | ENCSR000EFU |
| ESRRA | MA0592.3 | ENCSR000DYQ | ENCSR267NVP | ENCSR486IFJ |
| ETS1 | MA0098.3 | ENCSR000BKA | ENCSR672BSA | ENCSR000BKQ |
| ETV6 | MA0645.1 | ENCSR626VUC | ENCSR925TWM | ENCSR000FCE |
| FOXK2 | MA1103.2 | ENCSR861JUQ | ENCSR171FUX | ENCSR302AWT |
| GABPA | MA0062.3 | ENCSR331HPA | ENCSR000BJK | ENCSR290MUH |
| IKZF1 | MA1508.1 | ENCSR441VHN | ENCSR278JQG | ENCSR395HWC |
| JUND | MA0491.2 | ENCSR000DYS | ENCSR000EEI | ENCSR000EGN |
| MAFK | MA0496.3 | ENCSR000DYV | ENCSR000EEB | ENCSR000EGX |
| MAX | MA0058.3 | ENCSR000DZF | ENCSR000BTM | ENCSR000BLP |
| MAZ | MA1522.1 | ENCSR000DZA | ENCSR000EDN | ENCSR000EFX |
| MEF2A | MA0052.4 | ENCSR000BKB | ENCSR291MJH | ENCSR000BNV |
| MXI1 | MA1108.2 | ENCSR000DZI | ENCSR000EDU | ENCSR000EGZ |
| MYC | MA0147.3 | ENCSR000DKU | ENCSR000DLR | ENCSR000EGJ |
| NFATC3 | MA0625.1 | ENCSR437GBJ | ENCSR635SKD | ENCSR670FDA |
| NFIC | MA1527.1 | ENCSR000BRN | ENCSR000BQX | ENCSR796ITY |
| NFYA | MA0060.3 | ENCSR000DNN | ENCSR124APT | ENCSR000EGR |
| NFYB | MA0502.2 | ENCSR000DNM | ENCSR935GZV | ENCSR000EGQ |
| NR2C2 | MA1536.1 | ENCSR000EUL | ENCSR559ZKI | ENCSR000EWH |
| NR2F1 | MA0017.2 | ENCSR514VYD | ENCSR662EOU | ENCSR970NKQ |
| NRF1 | MA0506.1 | ENCSR000DZO | ENCSR000EEH | ENCSR494TDU |
| REST | MA0138.2 | ENCSR000BQS | ENCSR000BJL | ENCSR137ZMQ |
| RFX5 | MA0510.2 | ENCSR000DZW | ENCSR000EEA | ENCSR000EGO |
| SP1 | MA0079.4 | ENCSR000BHK | ENCSR334KIQ | ENCSR000BKO |
| SRF | MA0083.3 | ENCSR000BGE | ENCSR000BLV | ENCSR000BLK |
| TBP | MA0108.2 | ENCSR000DZZ | ENCSR000EEL | ENCSR000EHA |
| TCF3 | MA0522.3 | ENCSR000BQT | ENCSR911MML | ENCSR970OJY |
| TCF7 | MA0769.2 | ENCSR501DKS | ENCSR444LIN | ENCSR863KUB |
| USF1 | MA0093.3 | ENCSR000BGI | ENCSR000BGM | ENCSR000BKT |
| USF2 | MA0526.3 | ENCSR000DZU | ENCSR000EEF | ENCSR000EHG |
| YY1 | MA0095.2 | ENCSR000BNP | ENCSR000BNT | ENCSR000EWF |
| ZBTB33 | MA0527.1 | ENCSR000BHC | ENCSR000BHR | ENCSR876GXA |
| ZNF143 | MA0088.2 | ENCSR936XTK | ENCSR080UEM | ENCSR000EGP |
| ZNF24 | MA1124.1 | ENCSR072PWP | ENCSR791AGT | ENCSR099NCH |
| ZNF384 | MA1125.1 | ENCSR000DYP | ENCSR101FJU | ENCSR000EFP |
| ZSCAN29 | MA1602.1 | ENCSR412YGM | ENCSR422VAG | ENCSR635EXI |

